# Combined blockade of CXCR4 and PD-1 enhances intratumoral dendritic cell activation and immune responses against HCC

**DOI:** 10.1101/2023.11.06.564640

**Authors:** Satoru Morita, Kohei Shigeta, Pin-Ji Lei, Tatsuya Kobayashi, Hiroto Kikuchi, Aya Matsui, Peigen Huang, Mikael Pittet, Dan G. Duda

## Abstract

Immune checkpoint inhibitors (ICIs) have transformed systemic therapy for unresectable hepatocellular carcinoma (HCC). Nevertheless, their efficacy is limited to a small percentage of patients, leaving an opportunity for enhancement through synergistic combination therapies. We tested here the combined blockade of programmed death receptor 1 (PD-1) and CXCR4, a receptor for CXCL12 and a key mediator of immunosuppression in the tumor microenvironment in orthotopic grafted and autochthonous models of HCC. We evaluated tumor growth and survival outcomes and examined the underlying mechanisms using immunofluorescence, flow cytometry, RNA-sequencing, and transgenic mice experiments. Combined anti-CXCR4/PD-1 therapy had a robust impact on tumor growth and significantly prolonged survival in all murine preclinical models. The combination treatment successfully reprogrammed antigen-presenting cells, revealing the role of conventional type 1 dendritic cells (cDC1s) in the tumor microenvironment. Moreover, DC reprogramming enhanced anti-cancer immunity by facilitating CD8 T-cell accumulation and activation in the HCC tissue. The effectiveness of the anti-CXCR4 antibody/ICI combination treatment was compromised entirely in *Batf3*-KO mice deficient in cDC1 cells. Thus, combined ICI therapy with an anti-CXCR4 antibody has the potential to augment the anti-cancer effects and improve survival outcomes in HCC via reprogramming intra-tumoral cDC1 cells.

## Introduction

Liver cancer is the sixth most commonly diagnosed cancer and the third most common cause of cancer-related death^1^, with increasing incidence and mortality rates in the United States and Europe^2^. Hepatocellular carcinoma (HCC) represents the majority of liver cancer cases^3^. Surgery is a potentially curative treatment, but many patients are not eligible due to underlying liver disease or high tumor burden. As such, systemic drug therapy is vital in managing unresectable HCC. Sorafenib, a multiple kinase inhibitor (MKI), has been the first-line therapy for over a decade, followed more recently by other MKIs. Still, resistance to MKI monotherapy develops rapidly^4–6^. Combining immune checkpoint inhibitors (ICIs) and anti-VEGF inhibitors has emerged as a breakthrough therapy. Still, it requires further advances, with less than 30% response rates and high relapse rates among initial responders^7,8^. Therefore, new combination therapies to stimulate the durability of immune responses after ICIs are urgently needed.

Dendritic cells (DCs) can be categorized into two main subsets: conventional DCs (cDCs) for antigen presentation and plasmacytoid DCs (pDCs) for interferon (IFN) production. Among cDCs, conventional type 1 DCs (cDC1s) are potent antigen-presenting cells (APCs) and accelerate subsequent anti-cancer immune responses, while cDC2s stimulate CD4^+^ T-cell responses via MHC II^9,10^. Recent findings have highlighted that in addition to these, there is an entirely distinct subset of CCR7^+^DCs that partially share features with both cDC1 and cDC2^11–14^. Of these, cDC1s are essential in triggering CD8^+^ cytotoxic T lymphocyte (CTL) activation and tumor cell identification and destruction through antigen cross-presentation^15,16^. Defective antigen cross-presentation is a significant cause of cancer immunosurveillance failure^17^. Besides, accumulating evidence in recent years highlights the significance of cross-presenting cDC1 in harnessing the potential of ICB therapy^18–22^. The liver microenvironment has been found to induce a paucity of DCs in tumors compared to other organs^23^. The mechanisms that suppress cDC1 in HCC remain unknown, but inefficient distribution for encountering tumor antigens and insufficient crosstalk between DCs and CTLs may contribute to immune tolerance in HCC.

CXCL12 and its receptor CXCR4 are crucial for the interaction between cancer cells and their tumor microenvironment (TME). CXCR4 is present in various cell types, including immune, endothelial, and stem cells, with increased levels generally present in cancer cells^24,25^. CXCL12 is also expressed by multiple immune cells, endothelial cells, stromal fibroblasts, and stem cells, and cancer cells often produce CXCL12^26^. CXCR4 signaling activation has been linked with cancer treatment resistance, tumor growth, invasion and metastasis, myeloid-derived suppressor cells (MDSCs) recruitment, and angiogenesis^27–30^. CXCL12/CXCR4 axis inhibition is being tested in clinical trials for various cancers, including HCC, where CXCR4 expression levels are associated with tumorigenesis, progression, and poor outcomes^31–36^.

Previous reports have shown that the CXCL12/CXCR4 axis promotes fibrotic and immunosuppressive TME formation in HCC^37,38^. However, CXCR4 inhibition alone has largely been ineffective across multiple cancer models, suggesting that CXCR4 targeting should be tested in combination with other modalities^39^. Indeed, the effect of specifically blocking CXCR4 on standard anti-PD-1 immunotherapy in HCC remains unknown. Moreover, most previous studies and clinical trials tested AMD3100, a small molecule antagonist of CXCR4, which has a short half-life and a complex mechanism of action. Using antibodies with a long half-life and specificity will help elucidate the relationship between CXCL12/CXCR4 signaling and the HCC TME, reduce off-target effects, and improve effectiveness. In this study, we discovered the potential of anti-CXCR4 antibodies to reprogram the immunologically “cold” TME of HCC to an immunologically “hot” one, mediated by DCs, and the benefits when combining them with anti-PD1 antibodies, a current standard of care for HCC patients.

## Results

### Combined anti-CXCR4/anti-PD-1 antibody treatment induces significant anti-cancer effects and survival benefits in orthotopic models of HCC with liver damage

We first investigated the feasibility and effectiveness of combining anti-CXCR4 and anti-PD-1 treatments (CXCR4/PD-1) in established preclinical murine HCC models, including the orthotopically implanted RIL-175 (*p53/Hras* mutant) and HCA-1 HCCs with induced liver damage ^40,41^. Tumor growth was monitored by ultrasound imaging. After HCCs were established (i.e., reached 5-6 mm diameter), tumor-bearing mice were randomized to treatment with either an anti-PD-1 antibody (BioXcell) (10mg/kg i.p. every 3 days), an anti-CXCR4 antibody (10mg/kg i.p. every 3 days), combination of anti-PD-1 and anti-CXCR4 antibodies (10mg/kg i.p. every 3 days), or an isotype control IgG (10mg/kg, every three days) (**Fig.S1A**). We found that the combination therapy was feasible, with no significant toxicity. Moreover, the combination therapy group showed a significant reduction of tumor volume and an increase in median overall survival (OS) compared to all other treatment groups both in the anti-PD-1 therapy sensitive model (RIL-175 in C57Bl/6 mice) and the anti-PD-1 therapy-resistant model (HCA-1 in C3H mice) (**Fig. 1A, B** and **Fig.S1B, C**). Finally, we tested the feasibility and efficacy of dual CXCR4/PD-1 blockade in autochthonous murine HCCs induced using Cre-adenovirus injection in *Mst1*^-/-^*Mst2*^f/-^ mice with induced liver damage (**Fig.S2A**). The combination therapy resulted in a significant tumor growth suppression effect, as shown by ultrasound imaging and pathological evaluation of liver nodules (**Fig. 1C, D,** and **Fig.S2B**). These results demonstrate that anti-CXCR4/anti-PD-1 combination therapy is feasible in murine models of HCC with liver damage, which mimics the clinical presentation of some human HCC and provides superior outcomes compared to anti-PD-1 therapy alone.

**Fig. 1.**
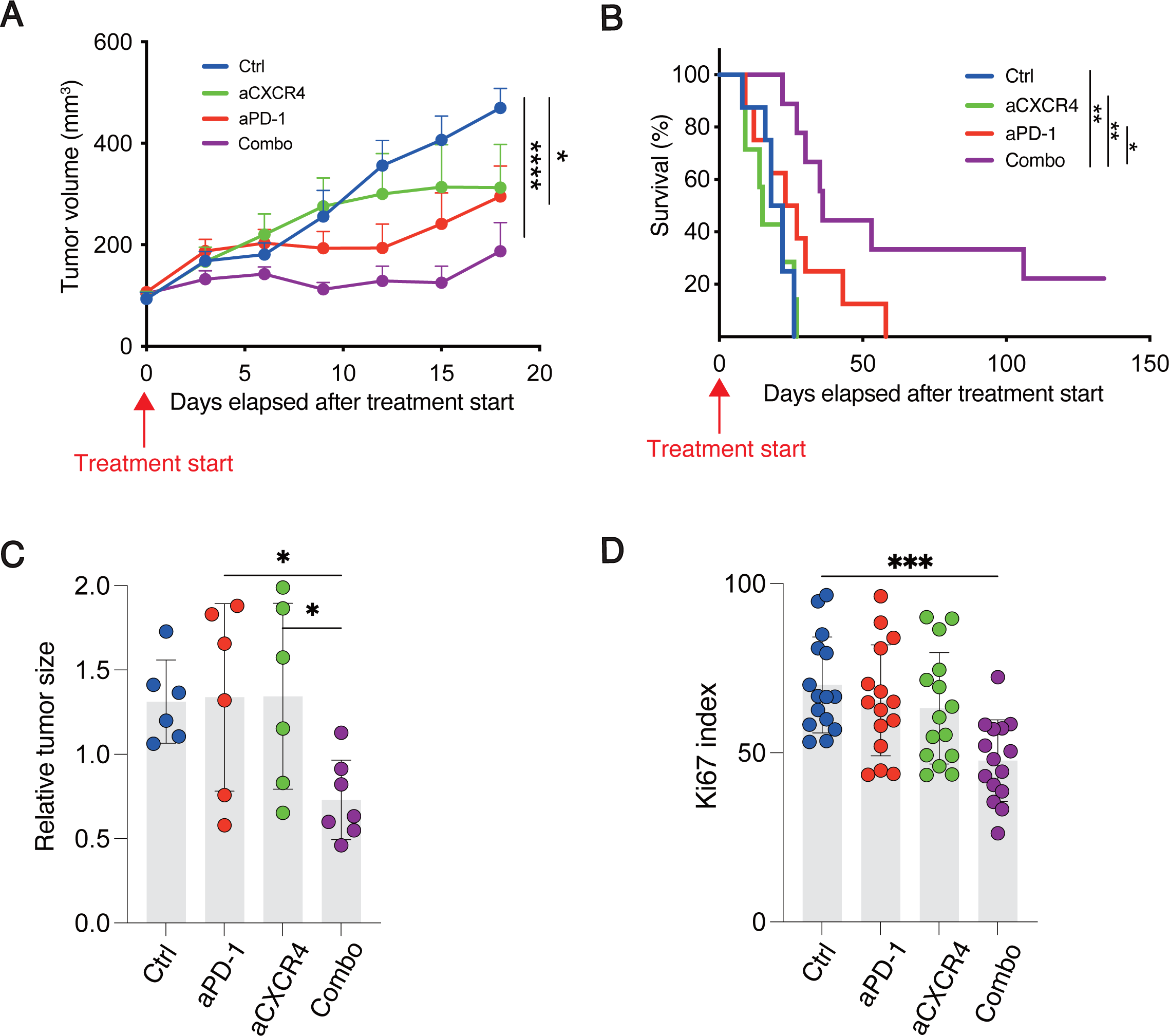
Combined anti-CXCR4/anti-PD-1 antibody treatment showed significant anti-cancer effects and survival benefits in orthotopic models of HCC with liver damage. (A-B) Tumor growth delay (A) and overall survival distributions (B) after a 3-week treatment with the anti-CXCR4 antibody, anti-PD1 antibody, their combination or IgG (isotype control) administered intraperitoneally at doses of 10mg/kg diluted in PBS thrice weekly in RIL-175 murine HCC orthotopic grafts in C57Bl/6 mice with CCl4-induced liver fibrosis. (C-D) Confirmation of potent anti-cancer effects for combination therapy in the autochthonous model (induced in *Mst1^-^*^/-^*Mst2*^F/-^ mice with underlying liver fibrosis) as measured by tumor volume using ultrasound (C) or tumor proliferation index (D)at day 14. Data are mean and s.d. (A, C, D) ***P < 0.001; **P < 0.01; *P < 0.05; two-way ANOVA (A) or log-rank test (B) or one-way ANOVA with Tukey’s test (C, D). N = 8 or 9 mice in all groups.

### Transcriptomic analysis indicates that combining CXCR4 and PD-1 blockade leads to an increase in antigen presentation and intratumoral dendritic cells (DCs) in murine HCC

We next repeated the experiment and collected tumor tissues from the treated mice bearing time-matched RIL-175 murine HCC. To understand the underlying mechanisms of how the combination therapy remodels the immune TME, we collected tumors from the RIL-175 models that received isotype control, single treatment, or combination treatment and performed RNA sequencing (RNA-Seq) analyses. The gene expression of the tumors in different groups was separated from each other by using principal component analysis (PCA), suggesting the remodeling of the TME after treatment (**Fig. 2A**). The PC1 contributes 90.3% of the variable and most genes in the PC1 were antigen processing and presentation associated genes. Next, we compared the differentially expressed genes (DEGs) between each treatment group and the isotype control group. We identified 100 DEGs that were only observed after combination therapy (**Fig. 2B**). Gene set enrichment analysis of these 100 DEGs showed a significant increase in pathways related to the T-cell receptor signaling pathway (**Fig. 2C**). Furthermore, T-cell activation-related signaling pathways were elevated in combination therapy group (**Fig. 2D**). Next, the expression patterns of genes related to DCs, which play a critical role for tumor antigen presentation, were compared among the groups. The combination therapy group exhibited higher expression levels of cDC1-related genes (**Fig. 2E**). Taken together, these data suggest that anti-CXCR4 and anti-PD-1 combination therapy enhances antigen presentation and DC frequency in the TME of HCC.

**Fig. 2.**
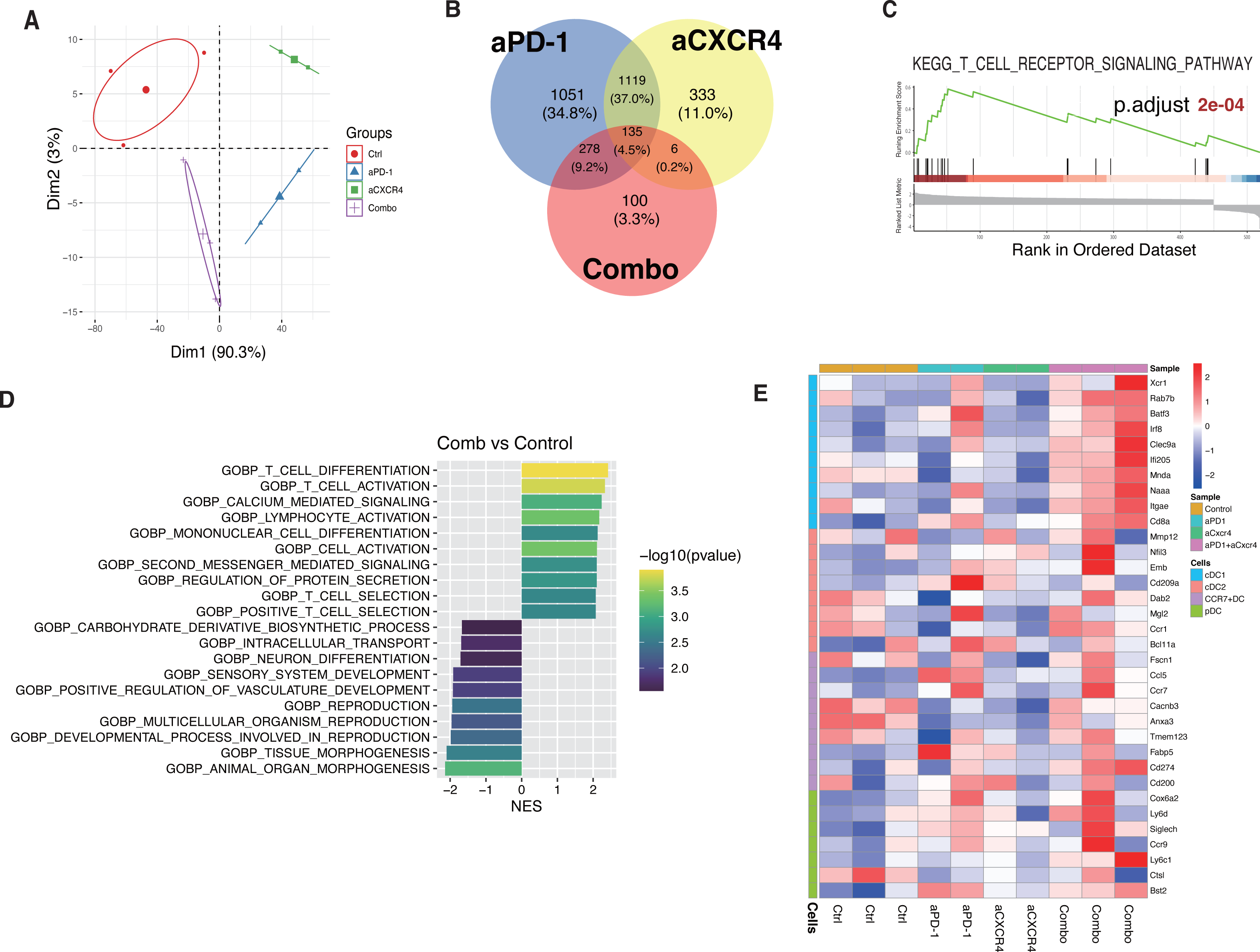
Transcriptomic analysis data from RIL-175 murine HCC after anti-CXCR4 therapy combined with anti-PD-1 blockade. (A) The principal component analysis (PCA) of gene expression profiles of murine HCC tissues from the control and treatment groups. (B) The Venn diagram represents the significantly differentially expressed genes between the treatment and control groups. (C) GSEA analysis showing T-cell receptor signaling pathway upregulated in combination therapy (D) Bar plot represents the top 10 most up- or down-regulated GO terms using Gene Set Enrichment Analysis. The GO terms are ranked by the Normalized Enrichment Score (NES). The color of the bar plot indicates the transformed P values. (E) The heatmap represents the gene expression of dendritic cell-related genes.

### Intratumoral conventional type 1 dendritic cells (cDC1) mediate the added benefit of CXCR4 blockade to anti-PD-1 therapy in HCC

To verify the results of transcriptomic data and examine the causal role of DC subsets, we first performed a flow cytometric analysis using time-matched tumor tissues from the four treatment groups. We found no significant difference in the groups’ overall frequency of cDC1s or cDC2s (**Fig.S3A, B**). However, when examining the distribution and expansion of DCs in TME by histological analysis, we found a significant increase in cDC1s in the center of tumors but not in the edge of the tumors after combination therapy, suggesting the combination therapy promotes the infiltration of cDC1. This effect was detected in orthotopic and autochthonous murine HCC models (**Fig. 3A-D** and **Fig.S4A**). The infiltration of cDC1 into the center of tumors may be due to active normalization of HCC vasculature, known to be induced by anti-PD-1 therapy^30^. Thus, we investigated the vessel normalization-associated gene signature in the bulk RNASeq data. However, we found no significant difference in vessel normalization signaling between the control and combination groups (**Fig.S4B**). This suggests that the added benefit of anti-CXCR4/anti-PD1 treatment is primarily cDC1-dependent.

**Fig. 3.**
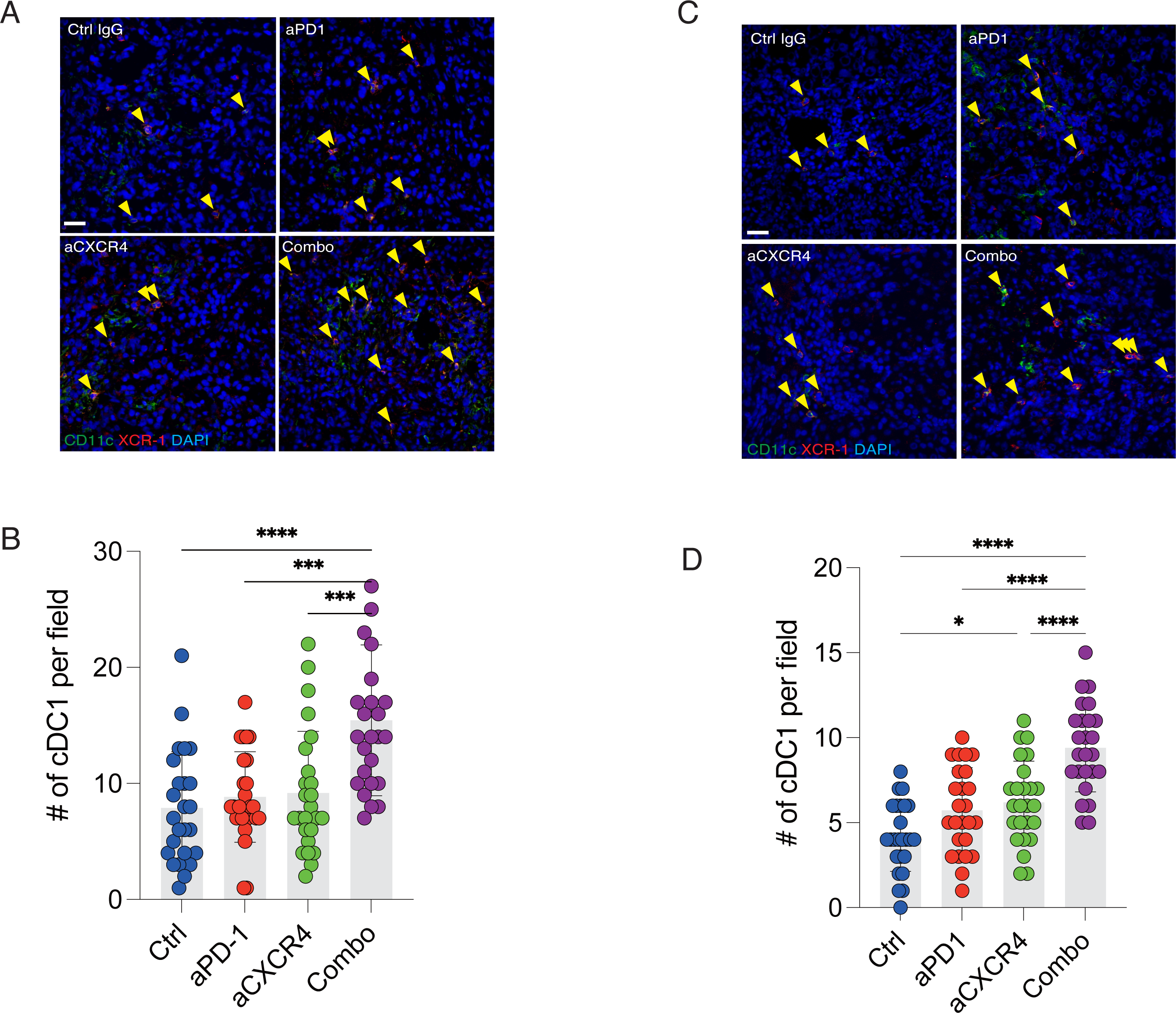
cDC1 infiltration in TME of HCC mediates the benefit of combined treatment with CXCR4 blockade to anti-PD-1 antibody therapy. (A-D) Combining anti-CXCR4 antibody with anti-PD-1 induces accumulation of cDC1 (defined as XCR1, CD11c double-positive cells) in both (A-B) orthotopic and (C, D) autochthonous model (arrowheads). Representative images (A, C) and quantification results (B, D) Data are mean and s.d. (B, D)***P < 0.001; **P < 0.01; *P < 0.05; one-way ANOVA with Tukey’s test (B, D) N = 7-8 mice in all groups. Scale bars, 50µm.

To determine whether cDC1s mediate the anti-cancer effect of the combined anti-CXCR4/PD-1 therapy, we next performed survival studies using *Batf3^-/-^/*C57Bl/6 mice, which specifically lack cDC1, and compared the results using C57Bl/6 WT mouse (**Fig.S5A**). As expected, murine HCC-bearing *Batf3^-/-^/*C57Bl/6 mice showed a deficiency of cDC1, as confirmed by flow cytometry (**Fig.S5B**). We found that the efficacy of the combination therapy (tumor growth delay and survival benefit) was abrogated entirely in the *Batf3^-/-^/*C57Bl/6 mice (**Fig. 4A, B** and **Fig.S5C-F**). These data demonstrate that the added anti-cancer effect and survival benefit induced by CXCR4 blockade, when combined with PD1 blockade, is mediated by cDC1.

**Fig. 4.**
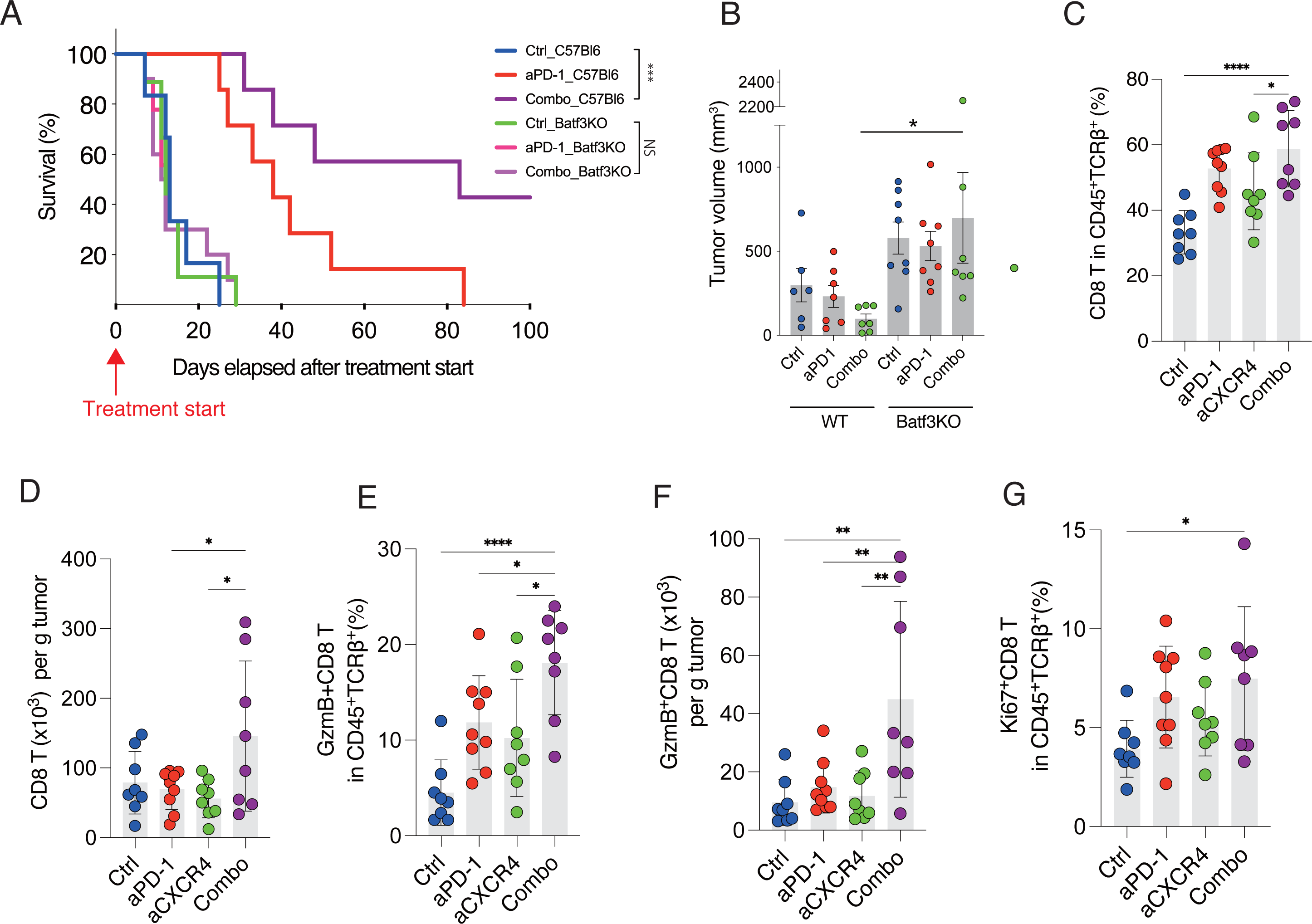
Combined treatment with anti-CXCR4 and anti-PD-1 antibodies promoted the activation and proliferation of cytotoxic T cells via cDC1. (A-B) The benefit of the combination treatment against murine HCC was compromised in the *Batf3*-KO (cDC1 deficient) mice. (A) Survival outcome, and (B) a comparison of tumor size at day 10 after starting treatment. (C-G) Combination therapy using anti-CXCR4 antibody and anti-PD-1 antibody induces proliferation and accumulation of activated cytotoxic T cells in TME of HCC: (C) frequency and (D) absolute numbers of tumor-infiltrated CD8 T cells; (E) frequency and (F) absolute number of GzmB expressing tumor-infiltrated CD8 T cells, and (G) Ki67 expressing proliferated CD8 T cells from the mice treated with control IgG, each single treatment or combination treatment. Data are mean and s.d. (B-G) ***P < 0.001; **P < 0.01; *P < 0.05; one-way ANOVA with Tukey’s test (A, B). N = 7-8 mice in all groups.

### Combined blockade of CXCR4 and PD-1 increases the activation of cytotoxic T cells in HCC

Tumor antigen presentation is mainly mediated by cross-presentation between cDC1 and CD8^+^ T cells, which leads to more functional CD8^+^ T cells and anti-cancer effects. Thus, we examined whether the combination therapy activates the cross-presentation of reprogrammed DCs and elicits CD8^+^ T cell activation in the TME of HCC by flow cytometry analysis using size-matched tumor tissues. The results demonstrated that the combination therapy significantly increased the CD8^+^ tumor infiltrated lymphocytes (TILs) (**Fig. 4C, D,** and **Fig.S6**). Tumors from the combination therapy group also showed a higher frequency of GrzmB^+^CD8^+^ and IFN-g^+^CD8^+^ T cells than either of the monotherapy groups (**Fig. 4E, F** and **Fig.S7A-E**). In addition, the combined blockade of CXCR4 and PD-1 increased the frequency of expanding Ki67^+^CD8^+^ T cells (**Fig. 4G**). These findings show that adding anti-CXCR4 antibody therapy to standard PD-1 blockade can increase the activation and expansion of intratumoral CD8^+^ T cells.

## Discussion

Although recent advancements in combination therapy using angiogenesis inhibitors and immunotherapy have surpassed sorafenib as the standard of care for HCC^7^, the therapeutic benefit remains limited, with almost all patients experiencing recurrences.

Several studies have investigated using CXCR4 signaling antagonists in treating solid tumors, including HCC. Despite this, these studies had significant limitations. One of the major limitations of CXCR4 antagonists for treating solid tumors is their lack of specificity and the short half-life of available drugs such as AMD3100^42,43^. To address these limitations, we evaluated the impact of combination therapy of anti-CXCR4 and anti-PD-1, a standard of care for HCC patients, in orthotopic and autochthonous preclinical HCC models. Our data suggest that anti-CXCR4 antibody enhances the anti-cancer effects of anti-PD-1 antibodies by increasing infiltration and accumulation of cDC1 and activated CD8 T cells in the TME of HCC. A recent study showed that CXCR4 inhibition with ICI enhances T cell retention in the TME, consistent with our results^44^.

cDC1 are well-known to play an essential role in anti-cancer immunity, but their role in the TME is not fully characterized and may depend on the organ microenvironment. Recent studies have demonstrated that cDC1s produce chemokines such as CXCL9/10, which enhances the intra-tumoral infiltration of effector T cells. This, in turn, can improve the efficacy of PD-1/PDL1 blockade^20,45^. In addition, we recently showed that recurrent (metastatic) HCCs lack PD-L1 expression and have poor antigen presentation function in patients^46^. This is consistent tith the findings that tumors growing in the liver tissue contain fewer DCs, partly responsible for the resistance to anti-PD-1 therapy in colorectal metastases or melanoma^23,47^. In addition to chemokine production, effective antigen presentation by DCs is crucial for activating CD8 T lymphocytes, which play a key role as effector cells in anti-cancer immunity. Thus, mediating DC accumulation and localization within the TME is of great interest therapeutically, particularly for anti-PD-1 therapy. The spatial distribution of immune cells within tumors is a critical aspect of the TME, and antigen-presenting cells exhibit intra-tumoral spatial heterogeneity in various cancers^48^. Here, we demonstrate that combination therapy with anti-CXCR4 and anti-PD-1 antibodies increased the infiltration of cDC1, especially inside the HCC lesions. Immune cell infiltration into the tumor center is critical for interacting with tumor cells. Indeed, we found that the combination therapy led to increased infiltration and activation of CD8 T cells and effective anti-cancer effects, which were completely compromised in mice lacking cDC1. In summary, our study demonstrates that combining anti-CXCR4 and anti-PD-1 antibody therapy is more effective than anti-PD-1 alone across murine HCC models and that the increased DC infiltration in the TME of HCC mediates the benefit. This effect is associated with increased infiltration and activation of CD8 T cells, a substantial anti-cancer effect, and prolonged survival. These findings may help improve the therapeutic efficiency of ICI-based therapy, especially against tumors growing in the liver, where DCs are scarce.

## Materials and Methods

### Cells and culture condition

Cell line authentication and mycoplasma contamination testing were performed before all experiments. RIL-175 cells were maintained in Dulbecco’s modified medium (DMEM) with 20% fetal bovine serum (FBS) with pyruvic acid and 1% penicillin/streptomycin. HCA-1 cells were maintained in DMEM with 10% FBS with pyruvic acid and 1% penicillin/ streptomycin.

### Mouse models orthotopic HCC and liver damage

To establish an orthotopic HCC mouse model with liver damage, RIL-175 cells were implanted in male C57Bl/6 mice and HCA-1 cells in male C3H mice, after 5 weeks of CCL4 treatment. To establish an autochthonous HCC mouse model with liver damage, we induced hepatocarcinogenesis by i.v. injection of a Cre-adenovirus in 4-week-old male *Mst1^-/-^Mst2*^f/-^ mice, and then induced liver fibrosis with CCl4 treatment for 12 weeks. When the largest tumors reached ∼5mm in diameter (after 12-16 weeks), mice were randomly assigned to a treatment group. Tumor growth was monitored by high-frequency ultrasound imaging. All experimental procedures were performed at least twice after obtaining institutional approval. Tumor growth was monitored using high-frequency ultrasonography every 3 days. Tumor volume was estimated using the following formula:

*Longest diameter x shortest diameter 2*^2^ */ 2*

### Treatments

Treatments were administered intraperitoneally twice weekly for 21 days at 10 mg/kg (anti-PD-1 antibody, anti-CXCR4 antibody, and immunoglobulin G [IgG] control). All animal experiments were performed after approval by the Institutional Animal Care and Use Committee of the Massachusetts General Hospital.

### Reagents

Anti-mouse CXCR4 and anti-mouse PD-1 antibodies, and isotype-matched rat IgG were provided by BMS per sponsored research agreement.

### Flow cytometry analysis

Tumor tissues were resected and minced, and fragments were incubated in Roswell Park Memorial Institute medium (RPMI) with collagenase, dispase, and DNase I for 30 minutes at 37°C. Digested tissues were passed through a 70-µm cell strainer and washed with HBSS/5mM EDTA. Single-cell suspensions were incubated with anti-mouse CD16/32 antibody (clone 93; Biolegend, San Diego, CA) before staining for immune cell markers for 15 minutes at room temperature. For the intracellular markers, cells were fixed and permeabilized with either the FoxP3/Transcription Factor Staining Buffer Set (eBioscience/Thermo Fisher Scientific, Waltham, MA) according to the manufacturer’s protocols. For cytokine staining, harvested cells were incubated in RPMI with a cell activation cocktail with brefeldin A (BioLegend) for 4 hours at 37°C, and stimulated cells were stained as described above. Monoclonal antibodies used for flow-cytometric analysis were CD45 (30-F11), TCRb (H57-597), CD8 (53-6.7), Ki67 (29F.1A12), IFN-γ (XMG1.2), GzmB (GB11), CD103 (2E7), XCR1 (ZET), MHC class II (M5/114.15.2), CD11b (M1/70), CD11c (N418), CD19 (6D5), Ly6G (1A8), NK1.1 (PK136), and Ly6C (HK1.4).

### Immunohistochemistry (IHC) and immunofluorescence (IF)

For IF on frozen sections of OCT-embedded tissues, each section was prepared at 8µm thickness for immunostaining. Sections were washed with PBS and blocked with 10% normal donkey serum for an hour at room temperature. Anti CD11c (CST #97585, 1:200) or XCR1 (Biolegend 109402, 1:200) antibodies were applied as primary antibodies overnight at 4°C, followed by the reaction with appropriate secondary antibodies (Jackson ImmunoResearch Laboratories) for 2hr at room temperature. All slides were mounted with ProLong™ Gold Antifade with DAPI (Thermo Fisher Scientific). Analysis was performed using a laser scanning confocal microscope (Olympus, FV-1000) in five random fields at ×200 magnification. Samples from orthotopic HCC were evaluated separately for tumor cores or edges, and samples from autochthonous HCC were evaluated randomly for each multicentric lesion. These data were analyzed with Fiji Is Just ImageJ (FIJI) and Photoshop (Adobe Systems Inc.). For IHC on paraffin sections, each section was prepared at 5µm thickness for immunostaining. Slides were deparaffinized in xylene for 10min, rehydrated through a graded alcohol series, placed in an endogenous peroxide blocker for 5min and washed with PBS. Sections were blocked with blocking solution (5% skim milk, 1%Triton X-100) for an hour at room temperature. Anti Ki-67 primary antibody (Millipore AB9260,) was applied overnight at 4°C, followed by peroxidase-conjugated anti-rabbit IgG as secondary antibody for 30min at room temperature. The sections were then developed with 3,3′-diaminobenzidine color solution for 3min at room temperature. Finally, hematoxylin was used as a chromogen and the slides were counterstained sequentially for 30s. Each slide was analyzed in five random fields under x200 magnification using bright field microscope (Olympus, BX40). The Ki-67 index of HCC growth (in the autochthonous model) was defined as the percentage of Ki-67 positive nuclei in each field and it was determined using ImmunoRatio plugins on FIJI by dividing the total intensity of positive nuclei by that of all nuclei in the field.

### RNA sequencing

Tumors were collected from mice treated with control IgG, anti-PD-1 antibody alone, anti-CXCR4 antibody alone, and anti-PD-1 antibody in combination with anti-CXCR4 antibody. Total RNA was extracted from the freshly isolated tumor tissues using Qiagen kits. Total RNA was extracted from freshly isolated tumor tissues using Qiagen kits. The RNA quality examination and sequencing library construction were conducted at the Molecular Biology Core Facilities of the Dana Farber Cancer Institute (Boston, USA). Raw sequencing data was subjected to quality control using FastQC. After quality control, Cutadapt was employed to remove low-quality bases and adaptor contaminations. The quality of the resulting clean data was examined again using FastQC software. Subsequently, Hisat2 was utilized to align the clean data to the mouse reference genome mm10, obtained from the Illumina iGenomes database. Following data mapping, SAM files, and BAM files were manipulated using samtools. The number of reads aligned to gene features was counted using HTSeq-count from the HTSeq package. Differentially expressed genes were identified by edgeR, using a cutoff of |log(fold change)| > 1 and a p-value < 0.01. GSEA analysis was performed using GSEA software with standard gene sets from the MSigDB database.

### Statistical analysis

Statistical analyses were performed using GraphPad Prism software (GraphPad Software, Inc.) or R software (for human adverse event analysis) and data was presented as mean ± SD. Specifically, the two-tailed unpaired Student’s t-test (parametric) was used for all comparisons between two groups, except for blood test comparisons, which were analyzed using the Mann– Whitney test (nonparametric). One-way analysis of variance (ANOVA) followed by Tukey’s post hoc test (parametric) was used for all comparisons between three or more groups. The Kaplan-Meier method was used to generate survival curves underlying the Log-Rank test and Cox proportional hazard model, and hazard ratio (HR), and 95% CI were calculated for statistical survival analyses for murine models. To analyze changes in body weight and tumor volume, we used two-way ANOVA with Bonferroni’s test (comparisons between two groups, parametric) or Tukey’s test (comparisons between three or more groups, parametric).

## Acknowledgments

The authors would like to express their sincere gratitude to T. Mempel for useful discussions, and M. Duquette, A. Khachatryan, H. Taniguchi, M. Galvan and S. Roberge (all MGH) for outstanding technical support

## Author Contributions

Conceptualization: SM, MP, DGD

Methodology: SM, KS, P-JL, TK, HK, AM, PH

Investigation: SM, KS, P-JL, TK, HK, AM

Visualization: SM, P-JL, TK, AM

Funding acquisition: DGD

Project administration: DGD Supervision: DGD

Writing – original draft: SM, DGD

Writing – review & editing: all authors

## Competing Interest Statement

DGD received consultant fees from Innocoll and research grants from Bayer, Exelixis, and Surface Oncology. No reagents from these companies were used in this study.

---

**Supplementary Figure 1. Anti-CXCR4 antibody combined with anti-PD-1 antibody treatment shows significant anti-cancer effects and survival benefits.** Timeline of tumor inoculation and treatments: After a 5-week course of oral 20% CCl_4_ to induce liver damage, tumors were inoculated orthotopically. Treatment was initiated 6 days after tumor implantation, and tumor-bearing mice were randomized to anti-PD-1 antibody (aPD-1) therapy, anti-CXCR4 antibody therapy, combination therapy with anti-PD-1 antibody and anti-CXCR4 antibody, or IgG (i.p.). Treatment was administered every three days and stopped at 3weeks after starting treatment (**A**). (**B-C**) Tumor growth delay (**B**) and overall survival distributions (**C**) after a 3-week treatment with the anti-CXCR4 antibody, anti-PD1 antibody, their combination or IgG (isotype control) administered intraperitoneally at doses of 10mg/kg diluted in PBS thrice weekly in murine HCA1 HCC orthotopic grafts in C3H mice with CCl4-induced liver fibrosis; n=12 mice per group. ***P < 0.001; **P < 0.01; *P < 0.05; two-way ANOVA with Tukey’s (**B**), log-rank test (**C**).

**Supplementary Figure 2. Efficacy of anti-CXCR4/PD1 combination therapy in an autochthonous model of HCC.** (**A**) Timeline of tumor inoculation and treatments: After injection of Cre-adenovirus in *Mst1*^-/-^*Mst2*^f/-^ mice, mice were administered CCl_4_ for 12 weeks to induce liver fibrosis. Tumor formation was confirmed by ultrasound imaging before the start of treatment. Mice were sacrificed to examine the effect 13 days after the start of the treatment. (**B**) The representative images of Ki67 immunohistochemistry; Ki67 index was used to quantify tumor burden in the *Mst1*^-/-^ *Mst2*^f/-^ autochthonous murine HCC model (n=7-8 mice per group).

**Supplementary Figure 3. Flow cytometry analysis of change in the overall frequency of dendritic cells among the groups.** Neither the overall frequency of cDC1 defined as XCR1^+^CD11b^-^ (**A**) nor cDC2 defined as XCR1^-^CD11b^+^ gated on CD45^+^Lin^-^MHCclassII^+^ CD11c^+^ (**B**) changed in response to anti-CXCR4/PD-1 combination therapy.

**Supplementary Figure 4. Anti-CXCR4/PD-1 combination therapy promotes the intratumoral distribution of dendritic cells (DCs).** (**A**) The quantification for the absolute number of cDC1s in the edge of the RIL-175 HCC tumor in C57Bl/6 mice. (**B**) GSEA analysis revealed no statistically significant disparity in vessel normalization signaling across the groups.

**Supplementary Figure 5. The efficacy of anti-CXCR4/PD1 combination therapy is compromised in *Batf3*-KO mice.** (**A**) Mice were randomized into three treatment groups: IgG control, anti-PD-1 antibody alone group, and the combination treatment with anti-CXCR4 antibody and anti-PD-1 antibody group. (**B**) Validation by flow cytometry plots that *Batf3*-KO mice lack cDC1s in the spleen. (**C-F**) The number of lung metastases (**C**), frequency of pleural effusion (**D**), ascites (**E**), and abdominal dissemination (**F**) in all treatment groups.

**Supplementary Figure 6. Gating strategy for CD8^+^ CTLs and their activation/proliferation status.**

**Supplementary Figure 7. Changes in CD8 T cell fractions in HCC after anti-CXCR4 antibody alone and anti-PD-1 antibody alone or in combination.** (**A, B**) Representative flow cytometry plots showing that combination treatment significantly increases the fraction of GzmB^+^CD8^+^ T cells (**A**), and Ki67^+^CD8^+^ T cells (**B**). Combination treatment increases the absolute number of IFNg^+^ CD8^+^ T cells (**C**) and IFNg^+^ GzmB^+^ CD8 T cells (**D**), and PD1^+^ CD8^+^ T cells (**E**) in the RIL-175 murine HCC model.

