## Supplementary figures and images for "Combined blockade of CXCR4 and PD-1 enhances intratumoral dendritic cell activation and immune responses against HCC"

### Supplemental Fig.1

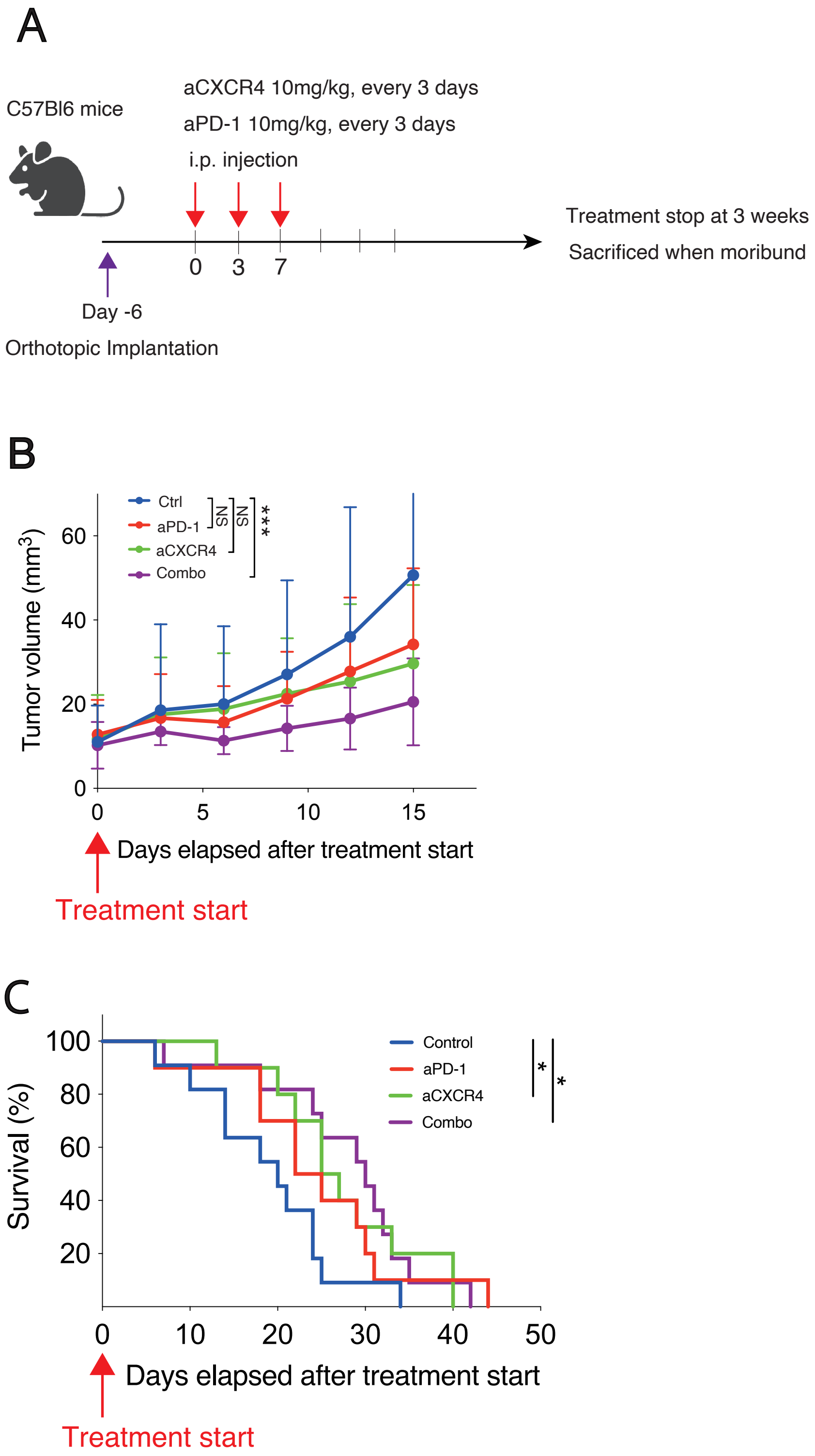

### Supplemental Fig.2

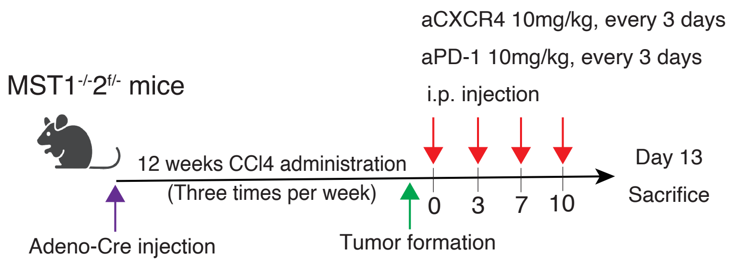

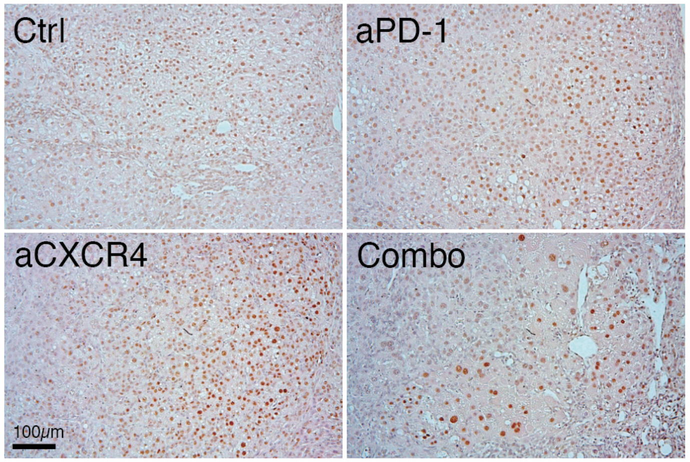

### Supplemental Fig.3

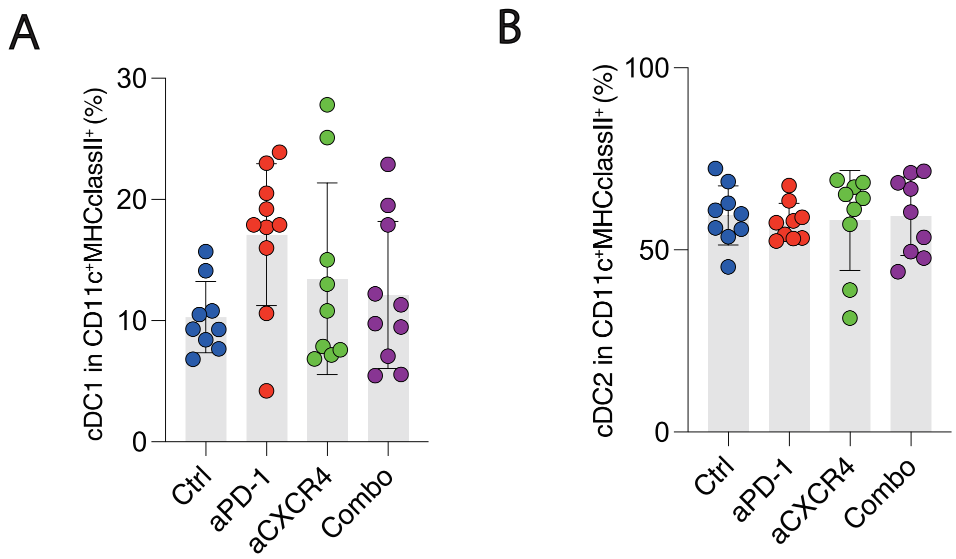

### Supplemental Fig.4

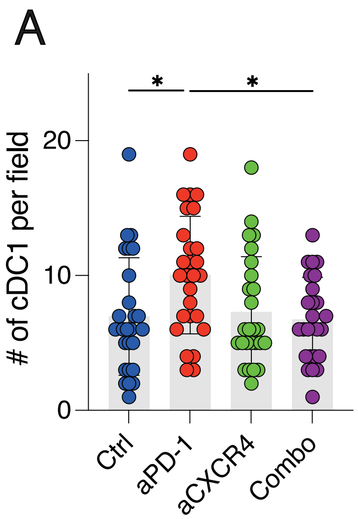

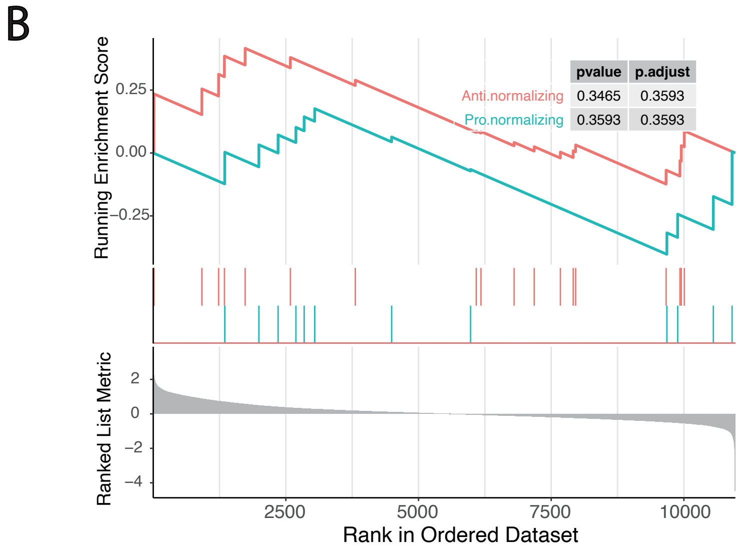

### Supplemental Fig.5

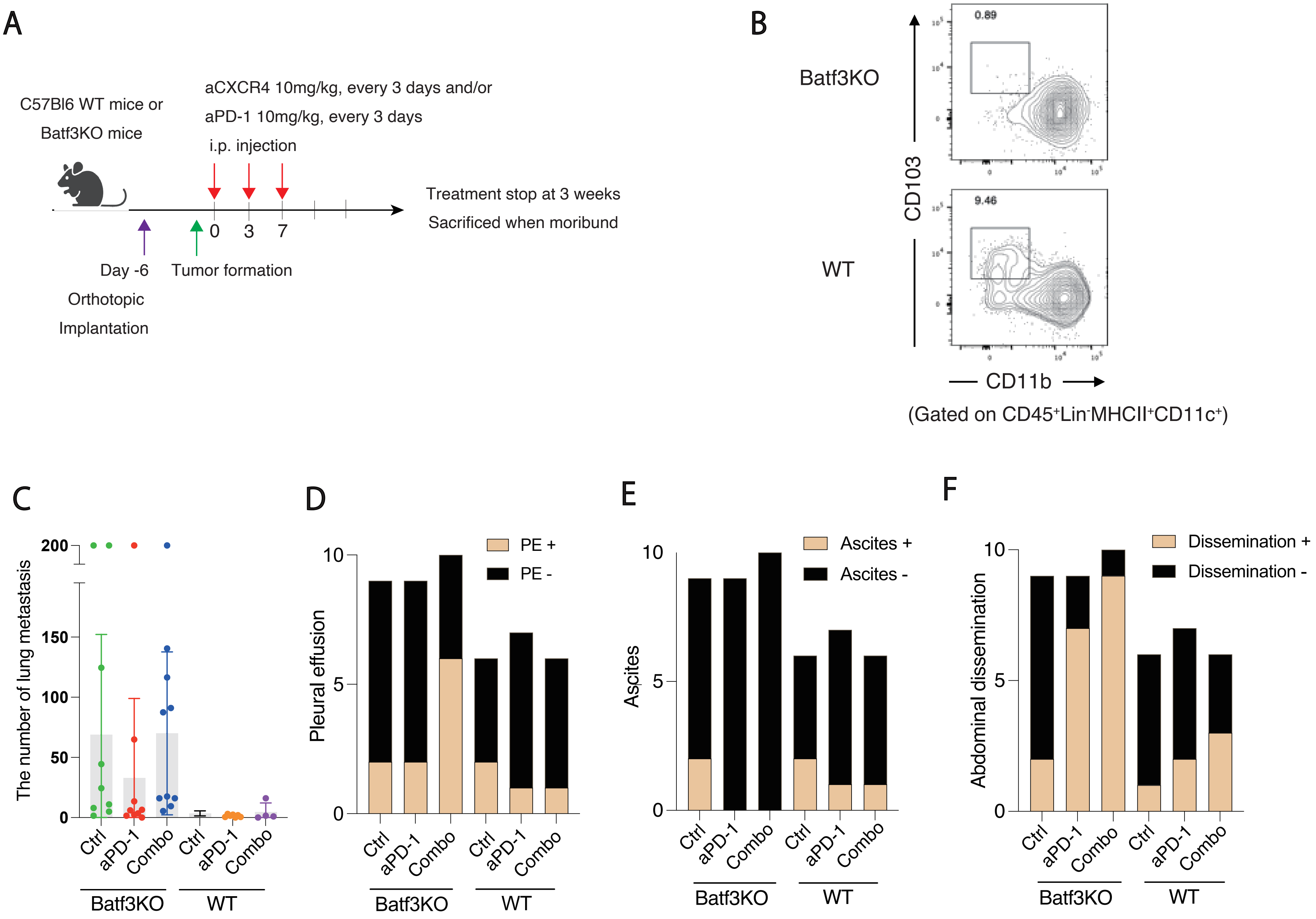

### Supplemental Fig.6

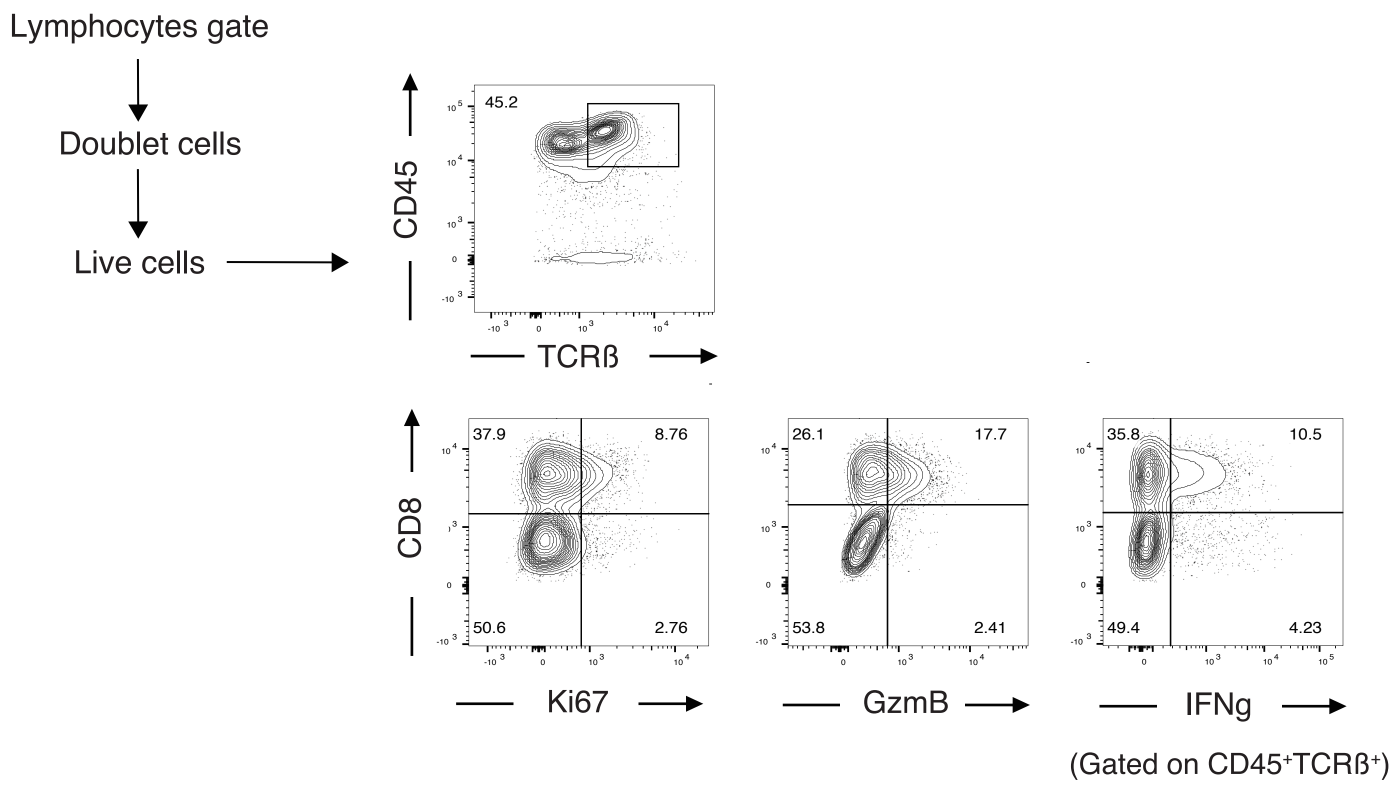

### Supplemental Fig.7

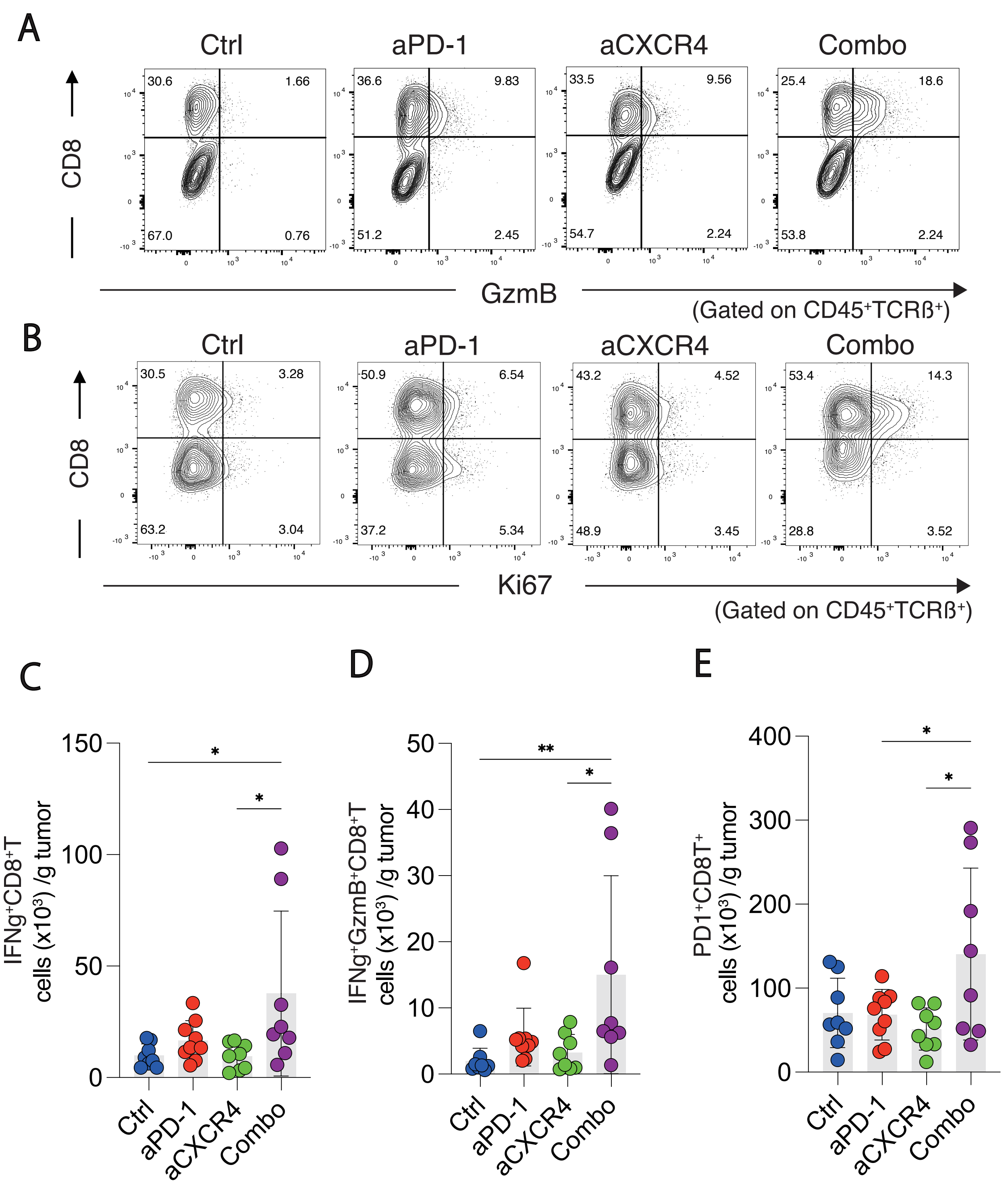
